# Decoding the Unintelligible: Neural Speech Tracking in Low Signal-to-Noise Ratios

**DOI:** 10.1101/2024.10.10.616521

**Authors:** Xiaomin He, Vinay S Raghavan, Nima Mesgarani

## Abstract

Understanding speech in noisy environments is challenging for both human listeners and speech technologies, with significant implications for hearing aid design and communication systems. Auditory attention decoding (AAD) aims to decode the attended talker from neural signals to enhance their speech and improve perception. However, whether this decoding remains reliable under severely degraded listening conditions remains unclear. In this study, we investigated selective neural tracking of the attended speaker under adverse listening conditions. Using EEG recordings in a multi-talker speech perception task with varying SNR, participants’ task performance—quantified through a repeated-word detection task—was analyzed as a proxy for perceptual accuracy and attentional focus, while neural responses were used to decode the attended talker. Despite substantial degradation in task performance, we found that neural tracking of attended speech persists, suggesting that the brain retains sufficient information for decoding. These findings demonstrate that even in highly challenging conditions, AAD remains feasible, offering a potential avenue for improving speech perception in brain-informed audio technologies, such as hearing aids, that leverage AAD to enhance listening experiences in real-world noisy environments.

## I. INTRODUCTION

Humans excel at selectively attending to a single speaker in noisy environments, a skill crucial for understanding speech in the real world. This ability is supported by the brain’s capacity to separate concurrent sounds and enhance the target speech [1], [2], [3]. Recent advances in neuroscience have demonstrated that neural tracking of speech features, such as the speech envelope, reflects this selective attention [4], [5], [6]. Through these neural responses, auditory attention decoding (AAD) has emerged as a promising technique, enabling the identification of the attended speaker based on brain activity in challenging multi-talker environments [7], which can be used to enhance that talker. In recent years, EEG-based AAD has shown substantial improvement in robustness and practical applicability, paving the way for real-world applications such as brain-controlled hearing devices and assistive technologies [7], [8], [9], [10], [11].

However, a key challenge in AAD is understanding whether it can remain effective under degraded listening conditions, particularly when the target speech is difficult to perceive. This raises a fundamental assumption: if speech is unintelligible, how can AAD reliably detect the attended talker to amplify their speech? In other words, AAD relies on neural tracking of the attended speaker, but neural tracking itself may depend on the intelligibility of the speech—posing a “chicken-and-egg” problem, where the two are interdependent. This question is especially critical for real-world applications, such as hearing aids, which are often needed most in noisy environments where the target speaker is difficult to discern. While previous studies have suggested that attention and intelligibility can influence the degree of neural tracking, the specific conditions under which perceptual breakdown renders AAD ineffective remain underexplored [12], [13], [14], [15].

In this study, we address this gap by investigating the robustness of neural tracking in situations where speech intelligibility is significantly compromised. Specifically, we designed a multi-talker speech perception experiment with varying levels of masker speaker and background noise, and measured participants’ speech perception using a repeated-word detection task [16], [17]. Simultaneously, we recorded EEG data to assess neural responses to both target and masker speech using canonical correlation analysis (CCA) [18], [19]. Our results reveal that selective neural tracking of the attended speaker persists even under highly degraded conditions, suggesting that the brain retains sufficient information to decode the attended speaker, regardless of significant intelligibility loss.

This work not only advances our understanding of the neural mechanisms underlying speech perception in noise but also provides new insights into the potential of AAD to improve listening in real-world noisy environments.

## II. METHODS

### A. Participants

Fourteen native American English speakers (7 males; mean ± standard deviation (SD) age: 25 ± 4 years) participated in the study. All participants reported normal hearing. They were compensated with a base payment, along with a performance-based bonus determined by task performance (1-back detection accuracy).

### B. Experiment

The study protocol was approved by the Institutional Review Board of Columbia University. Two loudspeakers were placed at ±45 degrees in front of the participants. Participants were instructed to focus on the target speech, with its gender and direction specified on a monitor. A non-target masker speech and stereo naturalistic background noises were played simultaneously (babble and pedestrian noise). All speech was in American English, synthesized by Google Text-to-Speech using four different voices (2 males and 2 females) to ensure a broad and consistent coverage across pitch and timbre. The target speech was fixed at 65 dB, with its SNRs ranging from −12 dB to 4 dB, adjusted by varying the loudness of both the masker speech and background noise.

To assist with target speech tracking in low-SNR conditions, each trial started with only the target speech with masker speech and background noise faded into the specified loudness after 3 seconds of the trial onset. These initial 3-second windows were excluded from the analysis. The experiment included 160 trials, each lasting approximately 35 seconds. During the task, participants pressed a buzzer when they detected repeated words in the target speech (mean interval between repeated words = 10s). After every round (16 trials), participants received real-time feedback on their task performance and were asked to briefly summarize the content, ensuring engagement.

### C. Preprocessing

For each of the 160 trials, 64-channel EEG data and buzzer responses were recorded using the g.HIAMP system (g.tec, Austria) and streamed through Simulink (MathWorks, MA, USA) at a sampling rate of 1200 Hz with a 60 Hz notch filter. EEG data were down-sampled to 100 Hz. Invalid channels with abnormal standard deviations were replaced using spherical interpolation of the remaining channels [20], [21], [22]. Speech envelopes for both target and masker speech were extracted using a nonlinear, iterative method [23] and down-sampled to 100 Hz to match the EEG recordings.

In this experiment, 1-back detection accuracy was used to quantify task performance during the experiment, serving as a proxy for perceptual accuracy and attentional focus as SNR varied. The average detection accuracy for each bin of SNR, ranging from −12 dB to 4 dB (bin size, 2dB), was fitted for each subject using *psignifit* toolbox [24], [25]. From the psychometric curves between SNR and task performance, we read the averaged task performance to reflect general speech perception under each SNR condition, as shown in Figure 2A.

**Fig. 1.**
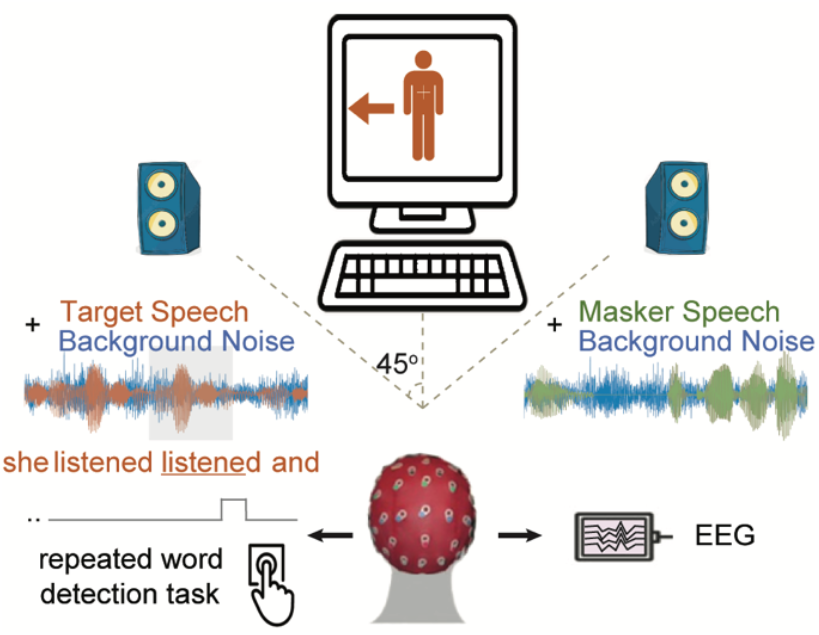
Experiment Schematic: Participants focused on target speech amidst background speech and noises.

**Fig. 2.**
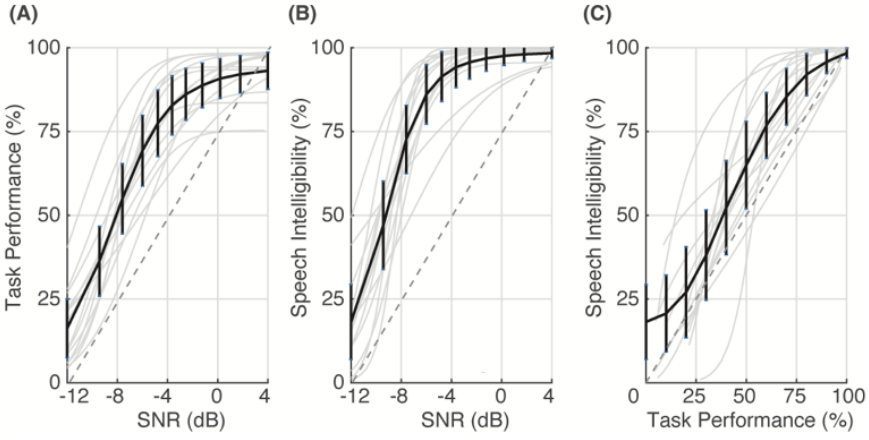
The relationships between task performance, speech intelligibility, and signal-to-noise ratio (SNR). While intelligibility is derived from a formal intelligibility test, task performance is based on the repeated word detection task during continuous speech, serving as a proxy for perceptual accuracy and attentional focus. The strong correlation between intelligibility and task performance supports the use of task performance as a measure of speech perception in our study,

Intelligibility, on the other hand, was measured separately before the experiment using the connected speech task [26]. Participants listened to short sentences at varying SNRs and repeated as many words as they could recall, allowing us to map SNR to intelligibility for each participant. Similarly, we fit psychometric curves for each subject by *psignifit* toolbox [24], [25], and they quantify the percentage of correctly repeated words at each SNR, as shown in Figure 2B. While task performance does not directly assess intelligibility, it correlates strongly with it, as shown in Figure 2C. Specifically, task performance reflects the participants’ ability to detect repeated words in continuous speech, providing a dynamic and real-time measure of their engagement and perception under varying SNR conditions.

### D. CCA Decoding Analysis

We used canonical correlation analysis (CCA) [18], [19] to evaluate the neural entrainment of speech envelope due to its simplicity and superior performance [19]. The CCA model estimates the relationship between stimuli (speech envelope) and neural responses (EEG) by transforming both into a shared space that maximizes their correlation. In this study, processed speech envelopes and EEG data were windowed with 200 ms and 400 ms overlapping receptive fields, respectively. The CCA model was trained using a leave-one-out cross-validation method for each participant.

For each participant and trial, the Pearson correlation between the speech envelope and EEG was computed to quantify neural speech tracking. The correlation for the target and masker speech is denoted as *r*_*T*_. and *r*_*M*_. The difference between *r*_*T*_. and *r*_*M*_., termed *Δ r*, was used to quantify selective neural speech entrainment, with *Δ r* > 0 indicating a larger neural tracking of the target talker relative to the masker talker and hence, a successfully decoded trial.

## III. RESULTS

To investigate how speech perception and neural speech tracking vary with signal-to-noise ratio (SNR), we analyzed the relationship between task performance, intelligibility, and SNR. As shown in Figure 2, task performance increases with improving SNR until reaching a ceiling at higher SNRs (Fig. 2A). This measure reflects participants’ ability to detect repeated words in continuous speech, showing higher accuracy at better SNRs. Intelligibility shows the same trend (Fig. 2B). Importantly, the strong correlation between task performance and intelligibility (Fig. 2C) validates the use of task performance as a reliable proxy for perceptual accuracy in the experimental conditions.

## A. Parameters influencing AAD performance

AAD decoding accuracy is defined as the percentage of trials where the target speech is successfully identified. In other words, if the correlation between a segment of the neural responses and the target speech envelope (*r*_*T*_) is greater than that of the masker speech (*r*_*M*_): *Δ r* = *r*_*T*_ − *r*_*M*_ > 0.

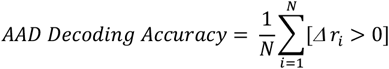

where *N* is the total number of trials.

As expected, both the duration and task performance are positively correlated with single-trial *Δ r* values and AAD decoding accuracy, as illustrated in Figure 3. The statistical significance of comparing *Δ r* with zero (Bonferroni-corrected t-test) is demonstrated in Figure 3B. Figure 3 shows that as the perception of the target speech decreases, the required decoding duration of the trial for successful neural decoding of the target speech also increases. However, neural decoding is still possible even when the target speech is barely detectable, as low as 25%, which is when only one out of four words is heard.

**Fig. 3.**
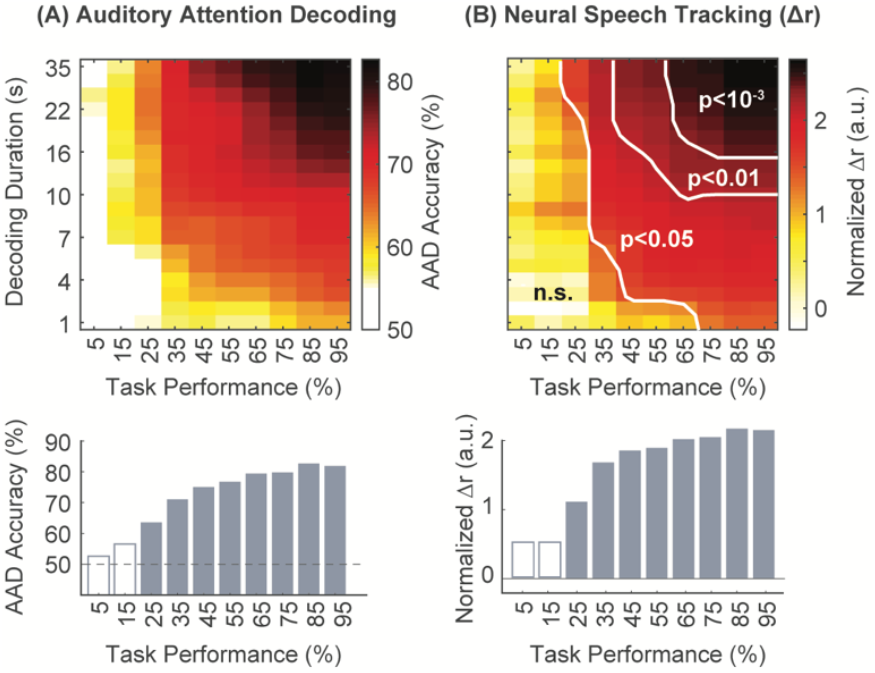
(A) Heatmap showing bootstrap-averaged AAD decoding accuracy across task performance and decoding durations. The bar plot represents AAD decoding accuracy at the maximum decoding duration of 35 seconds (full trial length), with white bars indicating non-significant results. (B) Heatmap showing bootstrap-averaged normalized t-stats for comparisons of Δr with zero (null), using a t-test with Bonferroni correction. The bar plot represents normalized t-values for task performance at a fixed decoding duration of 35 seconds, with white bars indicating non-significant results. For both (A) and (B), 300 bootstrap iterations with 30 samples per iteration were performed to prevent sample imbalance.

We conducted further analysis to isolate the effects of decoding duration and task performance on *Δ r*. As shown in Figure 4A, the mean of *Δ r* converges and remains constant after about 6 seconds, where more intelligible trials result in higher values of *Δ r*. Conversely, the standard deviation of *Δ r* is unaffected by task performance and is solely determined by decoding duration as shown in Figure 4B. In conclusion, increasing task performance improves *Δ r*, while increasing decoding duration reduces the statistical detectability of the single-trial AAD outcomes.

**Fig. 4.**
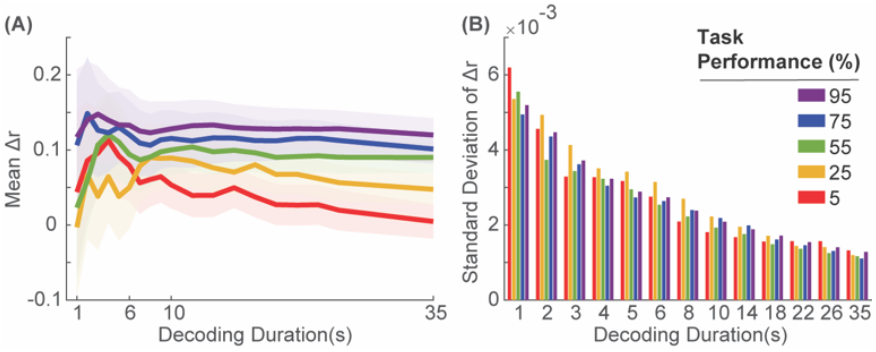
(A) Bootstrap-averaged mean *Δr* for different levels of task performance, across different decoding durations. (B) Bootstrap-averaged standard deviation of *Δr* for different levels of task performance. For both (A) and (B), 300 bootstrap iterations with 30 samples per time were used to alleviate sample imbalance.

### B. Simulation of AAD’s benefit

Since we have shown that AAD can be performed even in low intelligibility conditions, this suggests that AAD could be used to improve SNR in such scenarios, potentially leading to enhanced intelligibility. To explore this, we simulated the impact of AAD by comparing AAD-on and AAD-off conditions, as shown with red and black lines in Figure 5 accordingly. In AAD-on condition, we do the following:

**Fig. 5.**
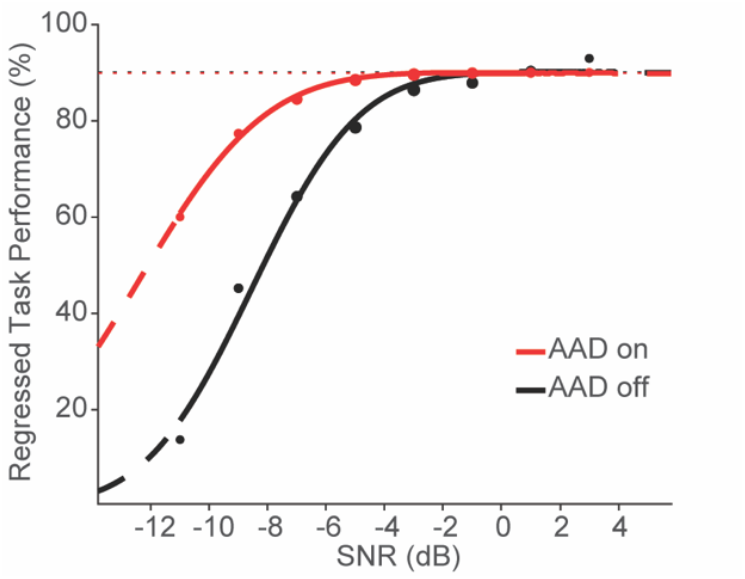
Simulation of the modulation effect on psychometric curves by implementing AAD (AAD-on, red line), compared to baseline (AAD-off, black line).

- When the predicted target speech is louder than the predicted masker speech, the overall loudness is adjusted to maintain 65 dB for the target, ensuring a consistent listening experience. No further amplification or noise reduction is applied to alter the relative amplitude of the target and masker speech, allowing for smooth attentional switches between speakers and minimizing detection errors.
- When the predicted target speech is quieter than the predicted masker speech, that predicted target speech is amplified to 65 dB, and the predicted masker speech is reduced to achieve a 0 dB SNR for the target. Specifically, under that SNR, the masker is still fully detectable.

In other words, the AAD-on condition operates on the left side of the psychometric curve and lifts up the task performance by increasing the SNR of the target speech. This simulation highlights the potential for AAD to significantly enhance speech perception in challenging listening conditions.

## IV. DISCUSSION

We investigated how the brain tracks attended speech in multi-talker environments with varying levels of background noise. Using a repeated-word detection task as a proxy for perceptual accuracy and attentional focus, which reflects intelligibility of continuous speech in real-time, we demonstrated that selective neural tracking of the attended speaker remains robust even when intelligibility is significantly degraded. This challenges the assumption that high intelligibility is necessary for effective neural decoding [27] and highlights the brain’s remarkable ability to retain sufficient information for decoding attended speech under adverse conditions. For example, neural tracking persisted even when only one out of every four words was perceived, extending the potential of AAD to low-intelligibility scenarios. Unlike traditional speech intelligibility measures, the repeated-word detection task captures a broader spectrum of perceptual and cognitive influences [28], [29], making it a dynamic and real-time tool for evaluating attention in noisy environments. These findings suggest a significant implication for AAD systems: they can function effectively even when the target speech is not fully intelligible, enabling the amplification and enhancement of speech perception in real-world noisy environments. This resilience positions AAD as a promising approach for brain-controlled hearing devices that improve communication in challenging auditory conditions.

Our findings also revealed a strong positive correlation between neural speech tracking and speech perception, aligning with prior studies [13], [16], [27], [30]. However, a key contribution of this study is the demonstration that neural tracking can be sustained even in environments with severely degraded intelligibility. This opens up new possibilities for applying AAD in real-world scenarios, where speech is often masked by noise. Additionally, we simulated the effect of AAD on speech perception and found that implementing AAD can significantly enhance perception, particularly in low-SNR conditions. These findings underscore the potential of AAD to optimize listening experiences in challenging environments. However, further research is needed to validate these findings in real-time, closed-loop systems and to explore the impact of cognitive factors such as attention and effort on decoding performance.

In future work, we plan to extend this research to more complex auditory scenes, where attended conversations involve turn-taking speakers who may also move in space rather than remain stationary [31]. We also aim to test these findings in individuals with hearing impairments to ensure the applicability of AAD in populations that stand to benefit most from enhanced speech perception technologies. Addressing these challenges will pave the way for developing adaptive, brain-informed technologies that enhance listening experiences in complex auditory environments.

